# Computational Finite Element Method (FEM) forward modeling workflow for transcranial Direct Current Stimulation (tDCS) current flow on MRI-derived head: Simpleware and COMSOL Multiphysics tutorial

**DOI:** 10.1101/704940

**Authors:** Ole Seibt, Dennis Truong, Niranjan Khadka, Yu Huang, Marom Bikson

## Abstract

Transcranial Direct Current Stimulation (tDCS) dose designs are often based on computational Finite Element Method (FEM) forward modeling studies. These FEM models educate researchers about the resulting current flow (intensity and pattern) and so the resulting neurophysiological and behavioral changes based on tDCS dose (mA), resistivity of head tissues (e.g. skin, skull, CSF, brain), and head anatomy. Moreover, model support optimization of montage to target specific brain regions. Computational models are thus an ancillary tool used to inform the design, set-up and programming of tDCS devices, and investigate the role of parameters such as electrode assembly, current directionality, and polarity of tDCS in optimizing therapeutic interventions. Computational FEM modeling pipeline of tDCS initiates with segmentation of an exemplary magnetic resonance imaging (MRI) scan of a template head into multiple tissue compartments to develop a higher resolution (< 1 mm) MRI derived FEM model using Simpleware ScanIP. Next, electrode assembly (anode and cathode of variant dimension) is positioned over the brain target and meshed at different mesh densities. Finally, a volumetric mesh of the head with electrodes is imported in COMSOL and a quasistatic approximation (stead-state solution method) is implemented with boundary conditions such as inward normal current density (anode), ground (cathode), and electrically insulating remaining boundaries. A successfully solved FEM model is used to visualize the model prediction via different plots (streamlines, volume plot, arrow plot).

## Introduction

Transcranial Direct Current Stimulation (tDCS) is considered a safe, portable, low cost, easy-to-use neuromodulatory technique that involves non-invasive delivery of weak direct current (typically 1 to 2.5 mA) to the brain. The resulting brain electric field (EF) modulates ongoing brain activity [1], influence synaptic efficacy [2], and produces plastic changes in excitability [3] and behavior [4]. Clinically applied tDCS protocols are generally designed and optimized in Finite Element Method (FEM) forward models [5]–[8]. Classical FEM forward models were developed for concentric spheres with biophysical volume conductor physics to examine the role of various dosage configurations [9]. tDCS dose is defined by electrode montage and current, while the resulting brain current flow is more complex and varies across individuals. Therefore, more recent forward models incorporated subject specific cephalic geometries, derived from Magnetic Resonance Imaging (MRI), to represent the human anatomy more accurately [10] with gyri/sulci precise resolution [5] depicting physiological detail in tissue micro-architecture through anisotropic tissue conductivities.

High resolution MRI-derived human head models are derived and segmented into tissue compartments with specific electrical properties. Electric current physics are applied to the volume conductor to investigate cortical EF distributions important in generating patient specific therapy protocols. This tutorial illustrates the pipeline of anatomically precise (voxel size of 1 × 1 × 1 mm) FEM forward modeling for individualized tDCS therapy dose design (Fig.1). Specifically, the guidelines include from data acquisition to solution generation in following steps: 1) Automated MRI scan segmentation; 2) Manual correction for tissue mask generation; 3) Electrode placement; 4) FEM mesh generation; and 4) Numerical FEM computation. The steps will be applied with tools such as Simpleware ScanIP (Synopsys, CA, USA) and COMSOL Multiphysics 5.1 (COMSOL Inc., MA, USA).

**Figure 1:**
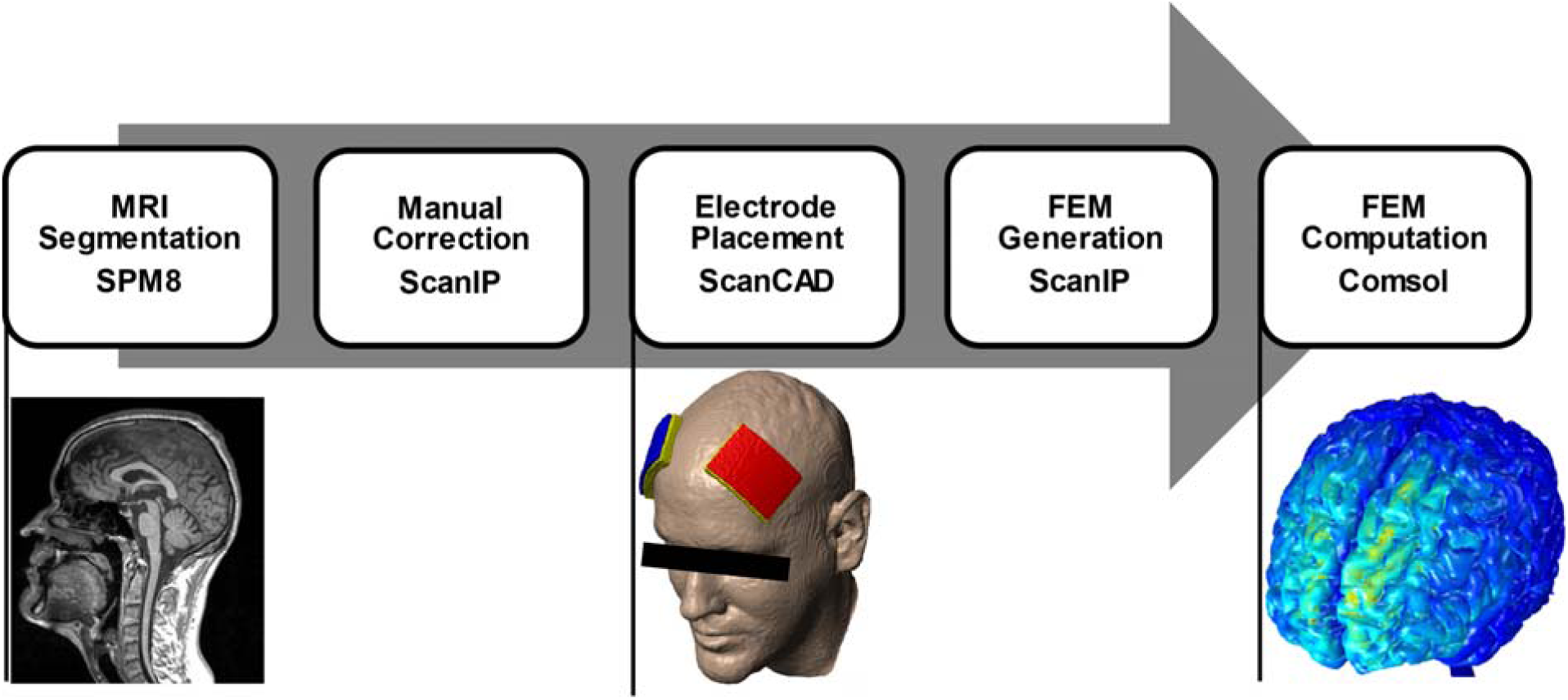
tDCS forward modeling pipeline. Including automated MRI segmentation, manual corrections, electrode placement, FEM generation and FEM computation (left to right).

## Materials and Methods

### MRI Segmentation (Automatic and Manual)

High-resolution T1-weighted magnetization prepared rapid gradient echo (MPRAGE) MRI scans from a 3-T Philips Achieva scanner (Philips Medical Systems, Best, The Netherlands) equipped with a Synergy-L Sensitivity Encoding (SENSE) head coil was performed on a healthy, neurological normal subject. Acquisition parameters were: TE = 3.2 ms; TR = 6.92 ms; flip angle = 8°; FOV = 256 mm; resolution = 256 × 256; slice thickness = 1 mm; no gap; and voxel size of 1 × 1 × 1 mm. The raw image data was then bias corrected and segmented into seven non-overlapping tissue compartments with an automated probabilistic segmentation routine from Statistical Parametric Mapping (SPM8, Welcome Trust Centre for Neuroimaging, London, UK). Additionally, an in-house MATLAB script (code available at: https://www.neuralengr.org/resources/) used to correct for automated segmentation errors [10].

The segmentation for current flow models of the head have several assumptions: continuity within tissues (laminar structure of skin, skull, CSF, grey matter) and continuity between tissues (i.e. no empty voxels). Manual segmentation enforces these assumptions and adds detail where the automated is insufficient. Typically, this occurs in bone due to its intricacy and lack of contrast within MRI scans. The folding of the cortex is often lost due to intricacy as well. With voxel-based segmentation, aliasing is a common problem within thin dynamic structures such as CSF and the sulci.

The output of the automated segmentation algorithm should start without overlap and without empty voxels. “Subtraction of masks on all slices” can remove overlapping tissues quickly if tissues priority is known. However, filling empty voxels is usually a messier process requiring multiple operations. To avoid empty voxels manual segmentation should aim to be zero sum, i.e. erasing any internal tissues should be followed by immediate filling with other tissues. To start, the segmentation should be rendered for an initial impression. Typically, manual segmentation can be divided into two primary steps: construction and detailing. Based on the initial rendering a decision can be made how much segmentation needs to be done. For noisy or irregular scans whole tissues may need to be reconstructed, which entails progressing slice-by-slice and tissue-by-tissue to trace or draw whole objects. Detailing is correcting holes, errors in continuity, and smoothing; which is independent of the bulk anatomical structure.

To reconstruct a complete head with little or no automated segmentation we work in layers. Working one tissue at a time allows completed tissues to act as stencils for neighboring tissues. Typically, we start with skull and move inward. The paint tool found under Image processing tab allows for tissue/material masks to be directly drawn on an image slice. Settings are available for brush shape and size as well as options for painting on ‘Active slice’, ‘Selection’, or ‘All slices’. Under most circumstances ‘Active slice’ should be selected. This restricts paint operations to the single active image slice rather than the entire volume. Left mouse bottom can be used to trace tissue of interest by navigating slice-by-slice with the paint tool. Right mouse button can be used to erase. If erasing, be aware that a void will be left behind within the head that needs to be filled. To streamline erasing and filling, it is often easier to work on a single boundary with a binary decision (i.e. the boundary of skull and skin or CSF and grey matter). Initially applying the union operation on skin with skull over all slices will allow skin to immediately fill any voids caused by erasing skull. Priority will be given to skull wherever the two tissues overlap, and skin will be subtracted from skull upon the completion of skull segmentation.

Whenever the identity of a tissue is unclear refer to an atlas or previously segmented models for reference. If the boundaries are not clear in a particular image, try to identify the tissue in subsequent slices and work back slice-by-slice to the problem area. The other orthogonal planes allow alternative views as well.

Because these scans are high resolution (1mm^3^ voxels), the segmentation should change only gradually between images. Shapes will have a spatial rate of change between slices that is typically conserved well beyond a single 1 mm thick slice. Sudden changes in segmentation between slices will lead to discontinuities and rough edges out of plane. Checking and/or correcting manually segmented tissues in the other orthogonal planes can help smooth shapes in three dimensions. The exception is when an object surface is in plane with the image orientation (i.e. occipital cortex in the coronal plane or cingulate gyrus in the sagittal plane). This can be leveraged whenever the segmentation is generally clean (i.e. little speckled noise within tissues and tissues are corregistered to the scan), but additional detail on a surface such as cortex or the skull needs to be quickly segmented. Segmenting surfaces in plane will reduce the number of slices needed to correct that surface and will in turn reduce the possibility of out of plane errors. For example, the temporal bone on the sides of the head are primarily in the sagittal plane. Manually segmenting the temporal bone in the sagittal plane will be faster than segmenting in the coronal plane – the former can be completed in roughly 20 slices of 1mm thickness, while the later spans over 100 slices.

After reconstructing the skull, the completed skull mask can be treated as a ground truth for subsequent operations. For example, the outer surface of CSF shares a boundary with skull. CSF can be quickly segmented while overlapping with skull, and then subtracted from skull to generate the exterior of CSF. CSF, in turn, forms the external boundary of grey matter. Segmenting CSF can be thought of as segmenting the grey matter surface. The segmentation of CSF is similar to that of skull, though, in some circumstances image contrast may allow for flood-filling of sulci when a clear distinction with grey matter is present (T2 sequence MRI’s in particular). The flood-fill operation fills areas within an image intensity range which is user defined. The specific range will vary between scans and within a scan itself, but generally T2 scans will have much brighter CSF than the surrounding skull and grey matter.

With the overall anatomy of the head segmented, a final “Detailing” check should be performed. Laminar tissues such as skin, skull, CSF, and grey matter should be rendered to check for errors in continuity, i.e. holes. Clicking on the 3-D view window, hovering over a point of interest (typically near a hole), and pressing the ‘p’ key on the keyboard will reposition the 2-D views to the point of interest. This procedure can be used to find isolated holes within a tissue. Holes are often clustered together, so a slice-by-slice analysis of the region is suggested. Breaks in continuity can be seen in 2-D as any point a laminar mask is not connected pixel side to pixel side. Note, that 3-D surfaces can have discontinuities out of view plane whenever continuous 2-D image slices are misaligned. This is easiest to spot by viewing all orthogonal views.

Other morphological filters such as the ‘close filter’ and ‘recursive Gaussian smoothing filter’ can be used in finishing segmentation. The close filter will collapse gaps within a mask. Generally, this is used on the air mask to reduce noise within the sinus cavities. The sinuses typically have poor contrast due to the combination of air and skull. To smooth and simplify the model, close filters applied to air will remove speckled noise by creating a solid enclosed volume of air without floating disjointed voxels of skull. Gaussian smoothing can be applied to the skin surface, while other tissues are often too intricate to smooth. Smoothing, however, can erode a mask and generate empty voxels. To only smooth the external surface and prevent empty voxels, first union skin with every other tissue mask in the head. Smoothing with a Gaussian sigma of near 1.5 mm is typically sufficient but will vary depending on the quality of the initial segmentation. Skin should then be subtracted from all other tissue masks.

Some steps for manually correcting or tracing tissue of interest:

1. Image processing tab of Simpleware has transform (resample, crop, rescale, padding, align, flip, shear, and shrink pad), segmentation (paint, unpaint, threshold, flood fill, 3D editing, etc.), morphological filters (erode, dilate, open, close, 3D wrap), smoothing (recursive gaussian, mean filter, media filter, etc.), and other additional tools (cavity fill, fill gaps, island removal, advance filters, etc.) to correct for non-uniformities.
2. Filling a gap or deleting a non-uniformity in any particular slice range can be performed by using a paint tool (most commonly used tool) found under segmentation. First step in using this paint tool is to specify the range of slices, and then appropriate brush type (square or disk) and size can be chosen to paint over the range of slices. Go to **Image Processing** → Select **Paint** → Choose **Shape and size of the brush** → Make **Slice selection** → Click **Apply**
3. Handling mask overlap is another import step in segmentation. Through simple boolean operation such as invert, union, subtract, and intersect, overlaps are generally handled. For example, if scalp spans over its actual thickness to skull, scalp can be subtracted from skull using “subtract” boolean operation so that the actual dimension of both scalp and skull are preserved. **Right click on Scalp** under **Masks** found under **Dataset browser** tree → Select **Boolean operation** and click on S**ubtract with** → Choose **Skull** → Select **Apply on all slices**

The generated tissue compartments represented gray matter, white matter, cerebral spinal fluid (CSF), air, skull, fatty tissue and skin/scalp.

### Electrode Placement

tDCS therapy dose is determined by 1) electrode montage, represented by electrode assembly properties such as selected materials and their geometry and the position on the scalp; and 2) the applied stimulation current (typically 1 to 2.5 mA). Electrode positioning is generally determined by International EEG 10-10 System [4] or by easy-to-use placement methods that aim to resemble EEG 10-10 scalp locations clinically [3]. The DLPFC can be targeted with a F_3_ (anode) - F_4_ (cathode) montage using 5 × 5 cm^2^ saline soaked sponge-pad electrodes with a current dose of 2mA (current density of 0.8 A/m^2^). CAD-Models (supported file formats: IGES, IGS STEP, STP, 3DS and STL) of sponge-pad or high-definition (HD) electrodes, created in SolidWorks 2013 - 2016 (Dassault Systems Corp., Waltham, MA), are imported and positioned on the scalp. The following steps will illustrate the positioning of a 25 cm^2^ sponge-pad electrodes are imported and positioned.

1. First step is to import the surface file (STL or CAD) of the electrode. Go to **Surface tools** → Click on **Import Surface** (CAD or STL) → Click **import**
2. Once imported, render fast preview to visualize the surfaces. Then using Positioning and Orientation feature, surfaces can be positioned at desired location. Specifically, this feature allows translation or rotation of the electrode to position them over the scalp. Go to **Surface tools** → Click on **Position and orientation** → Select **Surfaces** → Manually use the X, Y, Z **arrow or rotation icon** (Fig.3) placed over the electrode surface or insert **position (mm)/orientation (**^**0**^) to translate or rotate the electrode.
3. Next, the positioned surface is converted to mask. Go to **Surface tools** → Click on **Surface to mask** → Select **Surfaces** → Click **Generate mask**

**Figure 2:**
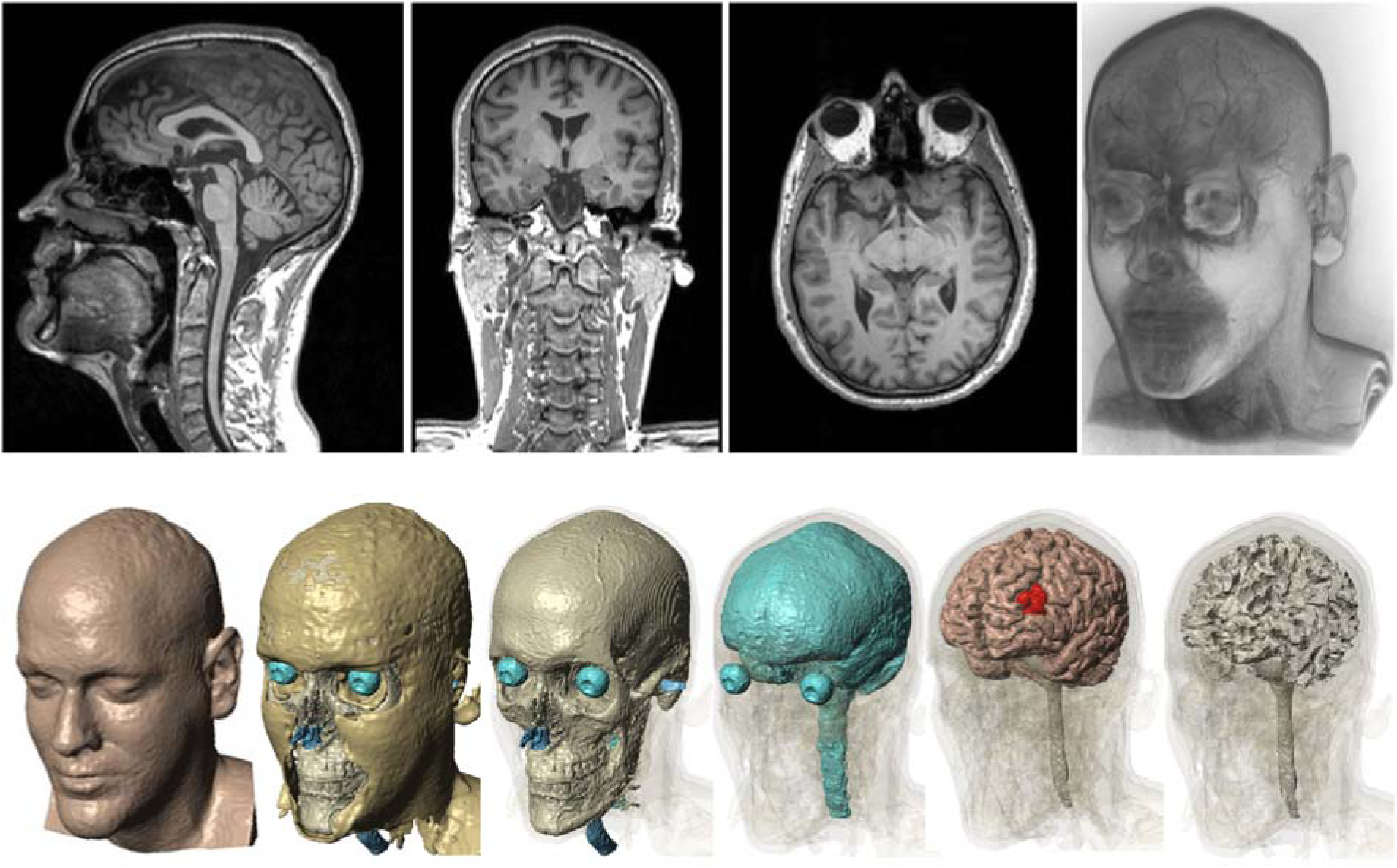
Segmented tissue compartments. representing skin, fat, skull, cerebral spinal fluid, gray- and white matter (left to right). The IDLPFC (red) was additionally segmented for later evaluation of anatomical targeting. Also, all models include air (blue) in the upper respiratory tract and the auditory channels.

**Figure 3:**
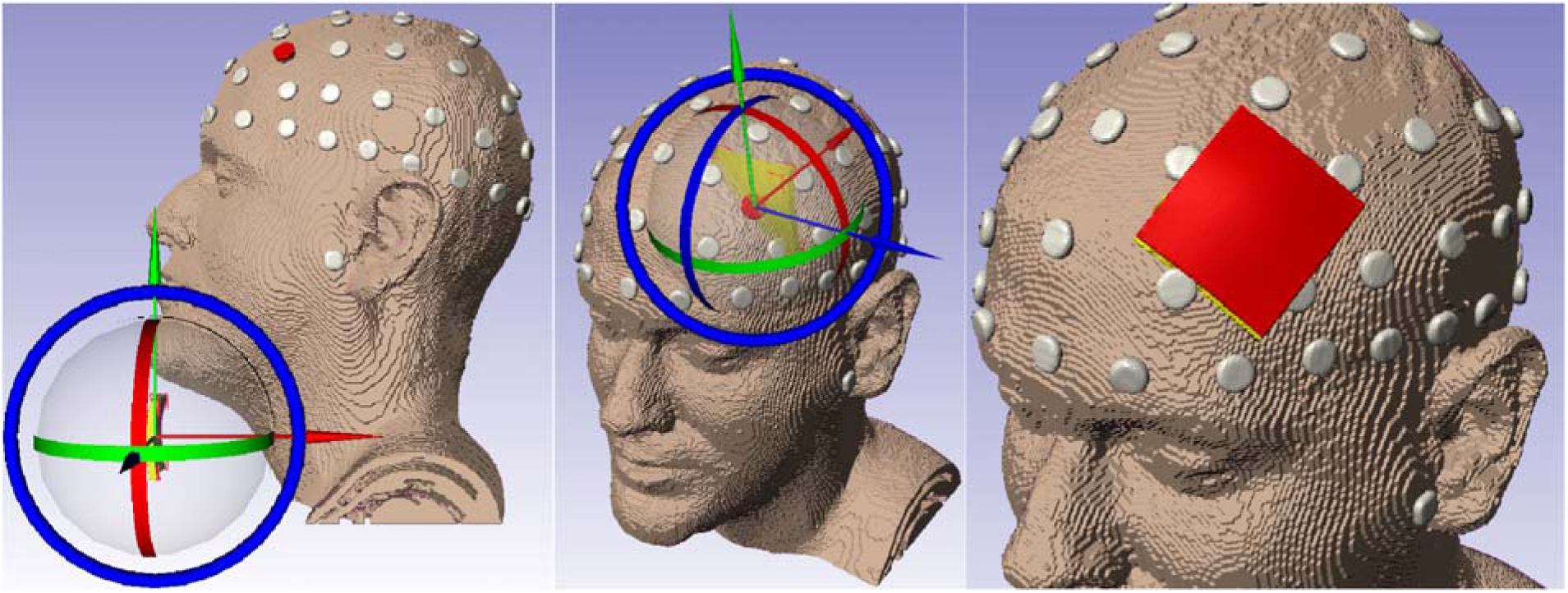
Deployed electrode positioning methods for an F3 anode. Bilateral encephalic electrode montage positioning of 5 × 5cm^2^ sponge-pads on EEG 10-10 scalp locations. With an F3 anode (red) and an F4 cathode (blue).

### FEM Generation

The FEM model implementation requires a three-dimensional (3D) volume mesh. Thus, the imported electrode montage and each of the eight segmented tissue compartments, including the lDLPFC as neuronal target, need to be meshed. Therefore, a total of 12 volumetric sub-domains consisting of approximately 10 × 10^6^ tetrahedral elements (Fig. 4) will be generated for later numerical computation. Simpleware ScanIP will be used to apply a Delaunay Triangulation Function with an adaptive (FE-Free) meshing algorithm.

**Figure 4:**
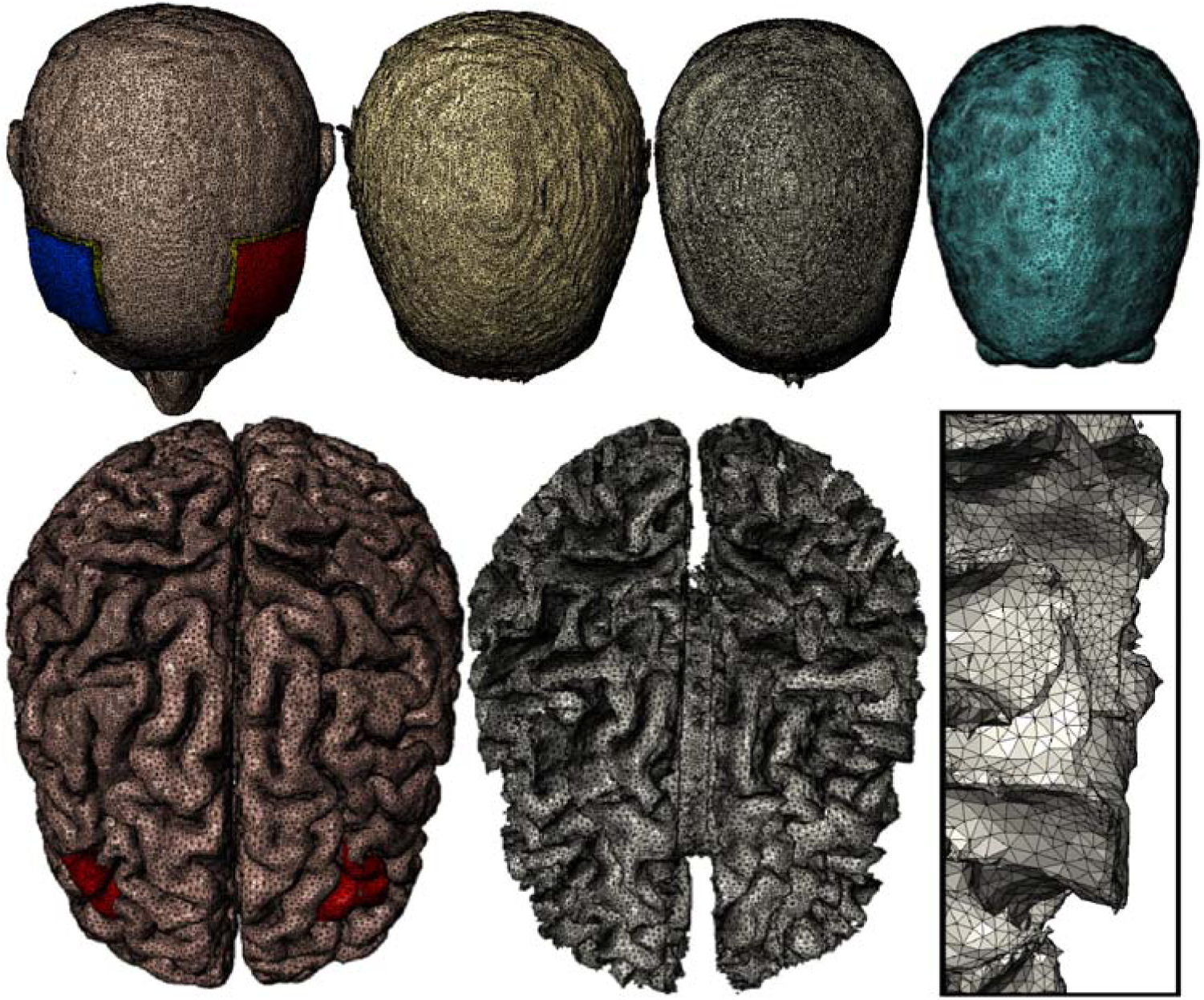
Exemplary volume mesh. Generated from seven tissue masks with and two sponge-pad 5 × 5cm^2^ sponge-pad electrode positioned over F3-F4. Top: depicting cathode (blue) and anode (red), saline soaked sponge (yellow), skin, fat, skull, air, and CSF (left to right). Button: illustrating gray- and white matter tissue compartment with a section view (left to right). The model entailed approximately 10.0 × 10^6^ tetrahedral elements with about 13.0 × 10^6^ degrees of freedom.

1. The most detailed geometries need to be meshed first to maintain realistic tissue architecture without losing detail through smoothing operations. The adaptive mesh algorithm will be applied to the mask in the following order: skull, air, skin/scalp, fat, CSF, DLPFC, gray matter, white matter, and sponge-. Therefore, the existing masks need to be arranged according to that hierarchy. Click Skull in the *data set browser* -> **Move up** -> Repeat with next mask
2. Next, a new model needs to be created from the existing masks of the working data set for the FEM meshing process Hold **Crtl** + each mask → **Right-click** on all marked masks → Add to active model. Simpleware will create a new model under Model (FE, active) in the bottom section of the *Data set browser*. Note that no EEG 10-10 marker will be needed in the FEM model.
3. Adjust model configurations for the mesh generation. **Right-click** on Model (FE, active) → **Model configurations** or go to **FE model** → **Setup model** In Model configuration window, go to Model Type → select **FE** Click on **Export Type** → select **COMSOL mesh volume (solid/shells) (v4.1 and later).** Selecting this option will output a COMSOL ready file with all domains selected according to the hierarchy of the mask in the Model builder. In **General** tab, select the **export length** unit to **Meters (m)** and **Coordinate system** as **Global**. The Options under **Smart mask smoothing** is left default. Go to Volume meshing tab → Click on **Mesh creation algorithm** -> Select **+FE Free** Click on the **Masks** under **Mesh density** and change the **Coarseness** as required. Normally, coarseness is set maximum for tissues with great details. For example, if coarseness is set as -10 for DLPFC, skull can have -16 coarseness. **Mesh refinement** tab can be used for refining certain boundaries of the tissues that needs special attention.
4. Create Mesh. Mesh generation takes approximately 3 hours with 24 Intel Dual Core processors, 3.1 GHz and 512 GB of RAM. Click on **Full model** under **FE Model** to initiate meshing
5. The resulting nodes and edges will be recorded as a COMSOL file (.mph). Export the mesh Go to **FE model** → **Export** → Select export directory → **Ok**

### FEM Computational Model and Solution Method

A 3D volume conductor model for electric currents with a stationary (steady-state) study type will be implemented using COMSOL Multiphysics 5.1. Subsequently, the generated mesh will be opened to assign isotropic tissue properties to each geometric entity (sub-domain), boundary conditions will be applied to the anode, cathode, and remaining external boundaries, and finally computed to predict the current flow.

1. Open exported COMSOL ready mesh file from the file directory
2. Apply **Physics** from the top navigation bar. Go to **Physics** → Select **Electric Currents (ec)**
3. Select Study Type Click on **Study** → **Add Study** → Select **Stationary** → Click **+ Add Study** icon
4. Next, an isotropic and homogeneous electrical conductivity value (in S/m) is assigned to each sub-domain of the head model. The adjusted values were as follows: air,1×10^-15^; skin, 0.465; fat, 0.025; skull, 0.01; CSF, 1.65; gray matter, 0.276; white matter, 0.126; electrode, 5.99×10^7^; saline-soaked sponge, 1.4; and conductive gel, 4.5 [12]. Since, the model is solved under quasi-static assumption, the relative permittivity value does not matter. So, a default 1 value is assigned with all tissue domains. Click on **Material** under **Components** in **Model Builder** → Click on each tissue (listed as a domain by default) and **assign electrical conductivity and permittivity** values as mentioned above.
5. Electrostatic volume conductor physics are then applied to the developed model. The following will be selected for each respective current density (*J*) with the corresponding surface normal (*n*). COMSOL’s AC/DC Module treats all exterior boundaries as electrically insulated (*n* * *J* = 0) and all interior boundaries as continuous across interfaces (*n* * (*J*_*in*_ − *J*_Out_) = 0). However, boundary conditions for anode and cathode must be selected manually.
  I. *Anode boundary condition* **Right-click Electric Currents (ec)** → Select **Normal Current Density** → Click on **Boundary Selection** → Set as **Manual** → **Click on exterior electrode boundary of the anode (F3)** in the *Graphics* window → Add to Selection (Plus sign) Click on Normal Current Density Type → **Inward current density** Measure the exterior electrode area (*A*_*Anode*_) to adjust the adjusted normal inward current density (*J*) according to 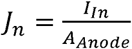 This corrects the boundary condition and compensates mesh artifacts on the electrode surface. Click on exterior anode surface -> Go to **Mesh** → Select **Measure** -> Calculated area is displayed in the *Messages* box at the bottom of the COMSOL window → Copy surface area from *Messages* → **Set Normal current density** *J*_*n*_ by dividing total current (2 mA) by the area as 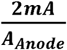 (approximately 0.8A/m^2^)
  II. *Cathode boundary condition* **Right-click Electric Current (ec)** → **Ground** → Click on **Boundary Selection** → Set as **Manual** → **Click on exterior electrode boundary of the cathode (F4)** in the *Graphics* window-> **Add to Selection** (Plus sign)
6. Click Compute to solve the model. The solver converges after approximately two hours. Click on Study 1 → Compute

## Results

A parental solution with the entire data set of the computed forward model allows separate or combined sub-domain, boundary and line-plots. Solution selections are typically created from a parental data set to analyze the created results in sub-sections. A surface plot of gray-matter is usually generated for the investigated head model (Fig. 5) to evaluate spatial EF intensities across the cortex depending on individual head anatomy and electrode positioning method. The sponge-pad electrodes are generally shown as wire-frame to reference the chosen montage on the skin. Radial electric field is depicted to differentiate inward (red) and outward current flow to locally interpret the resulting modulation as activation or inhibition. Also, vector field illustrations of current flow in combination with radial electric field distributions are created to depict directionality. Additionally, local electric field distribution illustrations of the targeted neuronal structure (here lDLPFC) are generated for comparison with the global EF distribution to evaluate stimulation focality.

**Figure 5:**
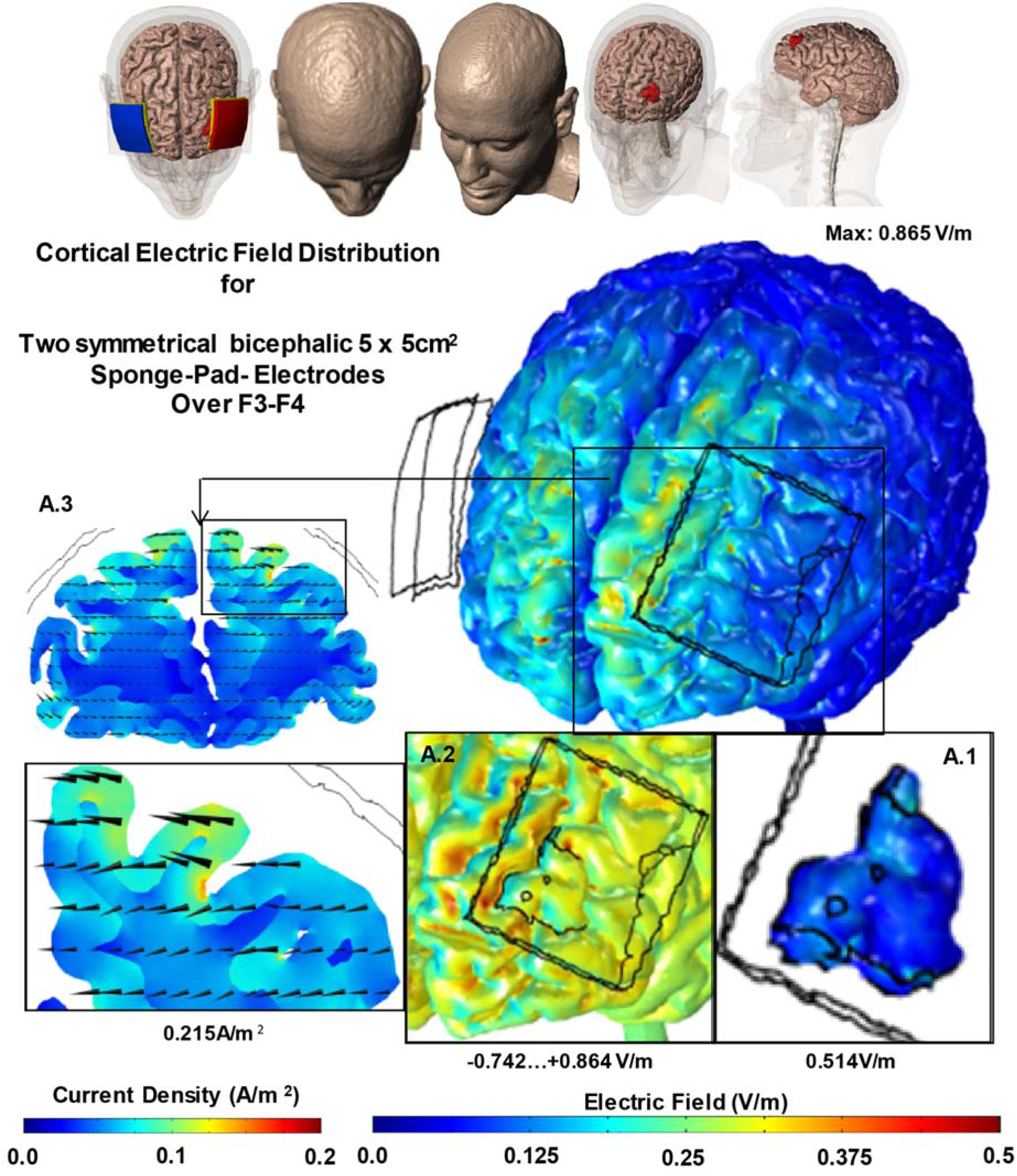
Cortical electric field distribution for two symmetrical bicephalic 5 × 5 cm^2^ sponge-pad electrodes placed over F3-F4. 2 mA, injected from the anode (red) to cathode (blue), resulted in the depicted electric field across the cortical surface. Cross-section views of the brain selected for maximal intensity (A.3), with magnified insets also showing the resulting vector field (cones), depict the produced current density distribution. The radial electric field across the lateral prefrontal region is also shown (A.2); scaled to ± 0.8V/m. The cortical electric field distribution in DLPFC (wireframe) is shown separately (A.1); scaled to 0.8V/m. Resulting peak intensities are depicted adjacent to each plot.

1. Uncheck automatic plot updates before a result selection is made to avoid frequent graphic updates. Click **Results** in Model Builder → **Uncheck Automatic update** of plots under Results Settings. A clear solution denomination simplifies later steps. Create separate selections from the parental data set by repeating the following steps for: gray-matter, lDLPFC, sponge-pad-electrodes and gray-, and white-matter. **Right-click on Data Sets** → Click **Add Solution** → **Right-click Solution 2** → **Rename** to Gray-Matter → **Right-click on Gray-Matter** → **Add Selection** → **Select Domain** from Geometric Entity Selection → **Set Selection to Manual** → **Select Gray-Matter** (Domain 7) in Selection List → Click **Add** to Selection (blue plus sign).
2. Individually allocate the data set selections to separate plot types. **Right-click on Electric Potential (ec)** under **Results** section in Model Builder → Select **Volume** → **Right-click on Volume 1** ≥ **Rename** to GM_electric Field.
3. Plot a false-color map in V/m of normal EF on the gray-matter volume. Select **GM_electric field** under **Electric Potential (ec)** → **Change the label** to GM_electric field → **Select Data set**: Gray-Matter. Go to expression (double headed green and red arrow) icon → **Select Electric Currents** > **Electric** > **Electric field norm (ec.normE)** or directly type the “**ec.normE**” in the expression window. The unit will automatically change to V/m. Click on Range and uncheck Manual color range, if checked as default. By default, the **Color table** is **Rainbow** which can be changed by clicking on the dropdown menu for color table. **Click Plot** at the top left corner to visualize the predicted electric field in gray-matter.
4. Cortical EF values in tDCS range from 0 to 2.5 V/m with peaks intensities in local “hotspots”. A range selection from 0 to 0.5 V/m allows sufficient contrast for result illustration. Note, that the range selection may vary depending on over-all head volume, montage selection including stimulation intensity or other factors. **Toggle Range** → Check: Manual color range → Set Minimum to 0 and Maximum to 0.5 V/m ≥ Click Plot (F8).
5. Add a wire-frame for the sponge-pad electrodes as a montage reference on the skin. **Right-click on Electric potentials (ec)** → **Choose Line** → **Select Data set**: Sponge-Pads in Domain settings window → **Replace expression**: **1** → **Toggle: Coloring and Style** → **Select Line type**: Line Coloring → **Select Coloring**: Uniform → **Select Color table**: Black → **Click Plot** (F8).
6. The local EF directionality may correspond to neuronal activation at locations of inward current flow and inhibition at outward current flow sights. Thus, this may be of significance when interpreting the investigating or designing a tDCS montage. Bidirectional radial EF plots demonstrate the distribution of cortical inward (red), and outward current flow (blue) in false-color maps. **Right-click Electric potentials (ec)** → **Surface** → **Right-click on Surface 1** → **Rename to Radial EF** → **Click on Radial EF in Model Builder** → **Select Data set**: Gray-Matter in Surface settings window → Under **expression** type: ***down(ec.Ex)*nx+down(ec.Ey)*ny+down(ec.Ez)*nz***. Note, that the directionality of a surface normal depends on the index of its domain. Consequently, the expression may need to be multiplied with -1. Set the range (step five above) to ± 0.5 v/m → Then **Click Plot** (F8).
7. Coronal section views (slice) through gray- and white-matter illustrate current flow patterns through the cortex. Coronal sections are generally selected for the coordinate of cortical peak CD. **Right-click on Electric Potential (ec)** → **Right-click on Slice 1** → Rename to Current Density (CD). **Click in Current Density (CD)** in Model Builder → **Select Data set**: Gray-Matter, White-Matter → replace **expression** (green and orange arrow) → Electric Currents ≥ Currents and charge > **Current density norm (ec.normJ**) or just type “ec.normJ” under expression → Toggle: **Plane Data** → **Select Plane**: yz-planes → **Entry method: Coordinates** → x-coordinate: 100 mm. Set the range (step five above) to 0 to 0.2 A/m^2^ → **Click Plot** (F8).
8. A vector field (cones) positioned over the cross-section of the CD plot additionally depicts current flow directionality. **Right-click on Electric Potential (ec)** → **Select Arrow Volume** → **Select Data set**: Gray-Matter, White-Matter → Under expression put current density (ec.normJ) → **Set x component** to 0 → **Toggle Arrow Positioning** → **Select Entry method**: **Coordinates under x grid points** → Select **Coordinate**: 100 mm → Select **Entry method:** Number of points under y and z grid points → **Toggle: Coloring and Style** → Select Arrow type: Cone → **Select Arrow length**: **Proportional** → **Select Arrow base: Tail** → **Set Scale factor to 50** → **Select Color Black** → **Click Plot** (F8).
9. Finally, a separate surface EF illustration of the targeted neuronal structure, here lDLPFC, shows its modulation in relation to the over-all cortical surface. **Right-click on Electric Potential (ec**) → **Volume** → **Rename to lDLPFC** → **Click on lDLPFC in Model Builder** → **Select Data set**: lDLPFC ≥ replace expression (green and orange arrow) to **ec.normE** → Select Color legend to include a unit scale for illustration → Set the range (step five above) to 0 to 0.5 V/m → **Click Plot** (F8).
10. Each previously generated plot may be disabled (F3) or enabled (F4) to view them separately or in combination. An example (Fig. 5) of a possible result arrangement is depicted below. Each plot is exported from the graphics window and saved as different image format. **Click Image Snapshot** in *Graphics window* → Select *Output Target*: File → **Output file format: .PNG** → **Specify filename and directory** → **Click Save**.

## Discussion

Computational FEM modeling studies are used to systematically optimize tDCS for clinical trials, experiments in cognitive function and/or therapy. A good understanding of the state-of-the-art modeling methods is needed for precise models to provide meaningful current-flow predictions. Our goal was to document and explain a widely used forward modeling technique while applying commonly used software packages to make the work-flow more broadly available. Here, we implemented an exemplary bicephalic sponge-pad head montage (F3 anode -F4 cathode) used for a common neuronal target (lDLPFC) to illustrate the necessary steps to predict current flow in the brain.

Forward modeling results are generally interpreted under the quasi-uniform assumption [11], however, note that the prediction of the clinical efficacy requires profound knowledge of the underlying brain functions. Furthermore, neuromodulation using tDCS requires additional therapy such as training to achieve optimal treatment outcomes. Computational modelling is the framework to rationally organize empirical data, formulate quantitative hypothesis, and test new interventions. Developing computational models requires the right balance of detailed multiscale model with appropriate reductionism [12]–[16]. A central motivation for modeling is that the interventional parameter space (dose, timing, task, inclusion citation, etc.) is too wide, given the cost and risk of human trials, for “blind” empirical optimization. Computational model is thus necessary for rational optimization of neuromodulation protocols [17], [18]. At early stages, such effort must be highly experimental data constrained [12], [19] and typically constrained to a limited range of dose settings. Computational FEM modeling is also the bridge by which data from animal studies can be rationally incorporated into models for interventions.

Computational models of current flow during tDCS in human models inform how the quasi-uniform assumption must be applied to support meaningful translational research. This assumption is based on a proportional relationship between neuronal excitation and the local electric field magnitude [20]–[23]. Certainly, the quasi-uniform electric field/current density representation is only an approximation for predicting the effects of tDCS, which is non-linear, time-dependent, and coupled system. However, the fact that it is nearly impossible to replicate tDCS induced electric field gradient across even a single hypothetical neuron between species -much less across the entire population of neurons - makes the quasi-uniform assumption a technical necessity in translational animal models.

## Acknowledgement

Source(s) of financial support: This study was partially funded by grants to MB from NIH (NIH-NINDS 1R01NS101362, NIH-NIMH 1R01MH111896, NIH-NCI U54CA137788/U54CA132378, and NIH-NIMH 1R01MH109289).

